# Mitochondrial Oxidative Phosphorylation Capacity in Skeletal Muscle Measured by Ultrafast Z-Spectroscopy (UFZ) MRI at 3T

**DOI:** 10.1101/2024.05.05.592570

**Authors:** Licheng Ju, Michael Schär, Kexin Wang, Anna Li, Yihan Wu, T. Jake Samuel, Sandeep Ganji, Peter C. M. van Zijl, Nirbhay N. Yadav, Robert G. Weiss, Jiadi Xu

**Author notes:** **Corresponding Author:** Jiadi Xu, Ph.D., Kennedy Krieger Institute, Johns Hopkins University School of Medicine, 707 N. Broadway, Baltimore, MD, 21205.

## Abstract

**Background:** To investigate the feasibility of rapid CEST MRI acquisition for evaluating oxidative phosphorylation (OXPHOS) in human skeletal muscle at 3 Tesla, utilizing ultrafast Z-spectroscopy (UFZ) MRI combined with the Polynomial and Lorentzian line-shape Fitting (PLOF) technique.

**Methods:** UFZ MRI on muscle was evaluated with turbo spin echo (TSE) and segmented 3D EPI readouts. Five healthy subjects performed in-magnet plantar flexion exercise (PFE) and subsequent changes of amide, phosphocreatine (PCr) and partial PCr mixed creatine (Cr^+^) CEST dynamic signals post-exercise were enabled by PLOF fitting. PCr/Cr CEST signal was further refined through pH correction by using the ratios between PCr/Cr and amide signals, named PCAR/CAR, respectively.

**Results:** UFZ MRI with TSE readout significantly reduces acquisition time, achieving a temporal resolution of <50 seconds for collecting high-resolution Z-spectra. Following PFE, the recovery/decay times (*τ*) for both PCr and Cr in the gastrocnemius muscle of the calf were notably longer when determined using PCr/Cr CEST compared to those after pH correction with amideCEST, namely 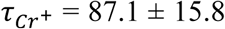 s and *τ*_*PCr*_ = 98.1 ± 20.4 s versus *τ*_*CAR*_ =36.4 ± 18.6 s and *τ*_*PCAR*_ = 43.0 ± 13.0 s, respectively. Literature values of *τ*_*PCr*_ obtained via ^31^P MRS closely resemble those obtained from pH-corrected PCr/Cr CEST signals.

**Conclusion:** The outcomes suggest potential of UFZ MRI as a robust tool for non-invasive assessment of mitochondrial function in skeletal muscles. pH correction is critical for the reliable OXPHOS measurement by CEST.

## Introduction

The creatine kinase reaction (CKR) and oxidative phosphorylation (OXPHOS) in muscle have previously been measured through changes in phosphocreatine (PCr) levels during physical activity and subsequent recovery phases with Phosphorus-31 Magnetic Resonance Spectroscopy (^31^P MRS) (1-9). Despite its utility, the adoption of ^31^P MRS in routine clinical settings is limited due to several impediments, such as its relatively low detection sensitivity when compared to conventional proton MRI, along with the need for specialized equipment, including ^31^P coils and concomitant transmit-receive hardware, which are not standard on clinical MRI systems. The Chemical Exchange Saturation Transfer (CEST) technique (10-15) has emerged as a promising approach for detecting low concentrations of Cr (16-20) and PCr (16,21-25) in tissues that does not require ^31^P system hardware. Nevertheless, isolating and quantifying PCr/Cr CEST signals from a crowded in vivo Z-spectrum poses a significant challenge. Initially, CEST experiments utilized a method called MTR_asym_ for contrast extraction, which calculates the asymmetry at a specific frequency by subtracting the signals equidistant from the water resonance. This method, also adapted for CrCEST, effectively reduced direct saturation effects (17,18,26). Despite this, MTRasym is unable to sufficiently negate other sources of interfering signals, such as those due to magnetization transfer contrast (MTC) and relayed Nuclear Overhauser Effects (rNOEs) (11,27) due to their asymmetry relative to water, as well as the influence of T_1_ relaxation times (28). To overcome these limitations, a new strategy, Polynomial and Lorentzian line-shape Fitting (PLOF), was introduced. PLOF enhances the specificity of Cr and PCr CEST signal extraction, especially in brain and muscle tissues at higher magnetic field strengths (22,23,29,30). Recent research indicates that the CrCEST exchange rate is approximately 270 s^-1^ in the brain (30-32) and less than 150 s^-1^ in skeletal muscle (33). Based on these findings, the PLOF method was applied to simultaneously extract PCr/CrCEST signals in skeletal muscle at 3T (33).

Nevertheless, there are two principal challenges for measuring CKR and OXPHOS in muscle using CEST MRI. Firstly, in healthy gastrocnemius muscles, PCr levels demonstrate an exponential recovery post-exercise, with an average recovery time of approximately 30-50 seconds (34-37). To capture the entire recovery curve accurately, the image acquisition must have a temporal resolution of 30-50 seconds, a criterion that presents significant challenges for traditional CEST MRI techniques. Secondly, intracellular pH levels in muscle tissue decrease during exercise due to two primary factors. The breakdown of ATP (adenosine triphosphate) releases hydrogen ions (H^+^), contributing to increased acidity. Moreover, intense exercise may cause the oxygen supply to the muscles to fall short of ATP demands. Under such conditions, muscles rely more heavily on anaerobic glycolysis, which leads to the production of lactic acid. CEST signals are highly sensitive to pH variations, potentially affecting the accuracy of PCr/Cr concentration measurements from their CEST signals. This study explores the feasibility to evaluate OXPHOS measured by post-exercise PCr recovery kinetics in human skeletal muscle at 3 Tesla using the ultrafast Z-spectroscopy (UFZ) MRI method (38-40). UFZ imaging significantly reduces the duration required to acquire a full Z-spectrum by spatially encoding the spectral dimension using a gradient applied simultaneously with the saturation pulse. Recently, this became more practical for brain application by aligning the slice direction with the spectral readout using so-called through-slice spectral encoding (TS) for UFZ (TS-UFZ). Consequently, amide, PCr, and Cr CEST maps can be generated in parallel using the PLOF Z-spectral data analysis. Our objective is to determine the capacity for mitochondrial OXPHOS in the skeletal muscle of healthy individuals by analyzing the dynamic PCr/Cr curves during in-magnet plantar flexion exercise (PFE). In addition, amideCEST imaging will be employed as a strategy to correct the PCr/Cr CEST signal measurements for pH influences.

## Methods

### MRI Experiments

All human studies conducted in this research were approved by the Johns Hopkins Medicine Institutional Review Board (IRB) and all participants (two male and three female healthy subjects, aged between 23 and 37 years old) provided written informed consent. A 3T Philips Achieva MRI system (Philips Healthcare, Best, The Netherlands) and a two-piece Sense XL Torso coil (Philips Healthcare) were used for all CEST experiments. The targeted calf was sandwiched by the two-piece coil.

The UFZ MRI sequence optimization was conducted on resting-state skeletal muscle. An optimum continuous wave saturation pulse of 0.6 µT and 2 s was applied according to previous study (33). Strength of the accompanying saturation gradient for the UFZ module was set to 1.0 mT/m in order to achieve a Z-spectral range of about 14 ppm (−4 to 10 ppm relative to the water frequency) across 40 mm. The FOV was 160 × 160 × 40 mm^3^ and acquisition voxel size was 5 × 5 × 0.4 mm^3^. Number of offsets in the Z-spectrum was 100. The center of the saturation pulse was set at 0, −3 and 3 ppm in the first three dynamic scans respectively to calibrate the saturation frequency in the UFZ spectra. In each dynamic, an unsaturated spectrum at 200 ppm was acquired as S_0_ prior to the saturated spectrum. Gradient echo (GRE) based segmented 3D EPI and spin echo (SE) based TSE readout were examined for comparison following the UFZ saturation module. For 3D EPI readout (41,42), UFZ MRI images were acquired with TFE factor 32 and turbo direction along y, EPI factor 17 along z, TR 46 ms and TE 22 ms with flip angle of 16°. The binomial 1-1 water excitation (WE) was selected for fat suppression. Total time for each dynamic was 10.4 s. The TSE readout parameters were TR = 4.055 s, TE = 8.3 ms, TSE factor = 64 with radial turbo direction, flip angle 90° and refocusing angle 120°, spectral presaturation with inversion recovery (SPIR) fat suppression was used. Total time for each dynamic was 24.3 s.

The PFE protocol included repetitively lifting a 16 lb weight for 2 inches at a rate of 1 Hz for 90 s. The CEST experiments were carried out throughout the process by alternatingly acquiring S_0_ (200 ppm) and S (3 ppm) UFZ MRI scans. With 26 dynamics, the total scan time was 10 mins 37 s under TSE readout. Each subject performed two PFE studies consequently, with a break of 10 minutes. A total of nine PFE experiments were recorded with five healthy subjects (one of the subjects performed one PFE).

Additionally, high-resolution T_2_ weighted images were collected for anatomical referencing using TSE sequence with a resolution of 0.44 × 0.44 × 4.0 mm^3^. T_1_ maps on human leg were acquired using the Dual Flip Angle (DFA) method as detailed previously (33).

### Data Analysis

All MRI images were analysed using MATLAB scripts developed in-house (The MathWorks, R2023a, Natick, MA). The Z-spectra were extracted using regions of interest (ROI) selected on T_1_ maps. Rician noise corrections were performed before further processing (32,33). A B_0_ correction was done using water saturation shift referencing (WASSR) with the assumption that the whole slice of tissue was homogeneous (43). The Z-spectrum was then linearly interpolated at 0.01 ppm per step. The extraction of the CEST signal was achieved by applying the PLOF method (28) over the Z-spectrum between 0.5 and 7.5 ppm. The ranges from 0.5 - 1.5 ppm and 5 - 7.5 ppm were selected for background fitting, while Lorentzian functions were assumed for the amideCEST, PCrCEST and partial PCr mixed CrCEST (Cr^+^CEST) peaks at 3.5 ppm, 2.6 ppm and 2 ppm, respectively. The extracted CEST signal ΔZ was obtained by taking the difference between the fitted background and the observed Z value.

To isolate mitochondrial OXPHOS activity in muscle tissue from the observed PCr/Cr CEST signals, two primary challenges must be addressed. Firstly, the CEST signal identified at 2 ppm represents a composite arising from both PCr amide proton (PCrCEST_2.0ppm_) and Cr guanidinium proton (CrCEST) signals. According to previous studies, the contribution of arginine guanidinium protons at 2 ppm from proteins in muscle is minimal (22), leading us to label this peak as to Cr^+^CEST in our analysis. The contribution of PCrCEST_2.0ppm_ to the Cr^+^CEST signal can be determined by multiplying the intensity of the PCrCEST peak at 2.6 ppm (PCrCEST_2.6ppm_) by a factor of 0.25. This factor 0.25 was derived by accounting for the fact that PCrCEST_2.0ppm_ involves one exchangeable amide proton, in contrast to PCrCEST_2.6ppm_, which involves two guanidinium protons. Furthermore, the exchange rate of the PCrCEST_2.6ppm_ protons is twice that of the PCrCEST_2.0ppm_ protons (21). Then, the pure CrCEST signal will be extracted following

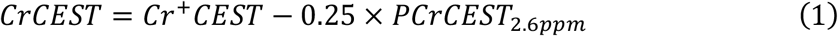

The second challenge involves addressing the pH influence on PCr/CrCEST quantification. To mitigate this, we can leverage the amide proton transfer signal at 3.5 ppm (44), dubbed amideCEST (centered at 3.5 ppm) here, as an indicator of *in vivo* pH. Assuming the exchange rates for amide and guanidinium protons are base catalyzed in the physiological pH range, the amide, PCr and Cr signals as a function of pH all follow (44-48) :

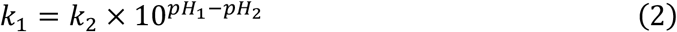

where k_1_ and k_2_ are exchange rates for the exchangeable protons at pH_1_ and pH_2_, respectively. In the slow exchange region, the CEST signal is proportional to the exchange rate (49). To derive PCr and Cr concentrations independent of pH variations, we normalize the PCr and Cr CEST signals against the amideCEST signal as follows:

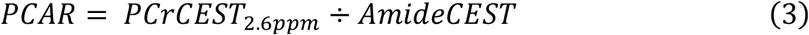

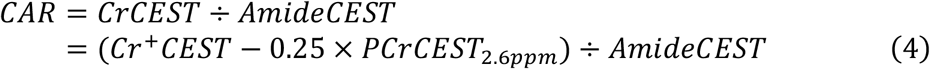

Here, PCAR represents the ratio of PCrCEST to amideCEST, and CAR denotes the CrCEST to amideCEST ratio. The PCr/Cr recovery/decay time constants, *τ*, following PFE are extracted using non-linear least-squares regression based on the following equation.

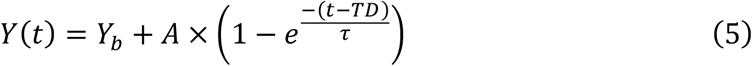

In which, *Y*(*t*) represents PCr or Cr signal intensity at any time (t); *Y*_*b*_ is the baseline Y value and A is the amplitude change factor in *Y* from the baseline value; TD is the time delay.

## Results

### UFZ sequence optimization

Typical UFZ spectra are shown in Fig. 1A. For the UFZ experiments, the saturation offsets need to be calibrated as the x-axis in the UFZ spectrum is the sub-slice number. Two sets of dynamic scans (offsets Δ*ω*: [200, −3] ppm, [200, 3] ppm) needed for the offset calibration were acquired before the PFE study under identical gradient and FOV specifications.

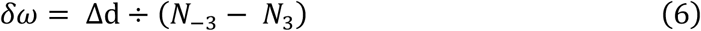

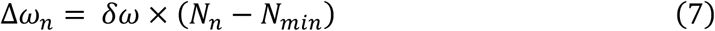

in which, N_-3_ and N_3_ are the sub-slice numbers with the minimum of the Z-spectrum centered at −3 and 3 ppm, respectively. (N_-3_ - N_3_) corresponds to the offset difference Δd in Fig. 1A, *δω* is the Z-spectral resolution. N_n_ and N_min_ are the sub-slice numbers of one point of interest on the UFZ spectrum and the sub-slice with minimum Z value along this Z-spectrum, respectively. Δ*ω*_*n*_ is the offset value in ppm for this interested point. Through the calculation above, typical Z-spectra were obtained and presented in Fig. 1B for UFZ with different readout schemes. For UFZ employing 3D EPI readout, the Z-value at 0 ppm (0.23) was significantly higher compared to that recorded with TSE (0.035). Figures 1C and 1D illustrate the extraction and quantification of CEST contrasts using the PLOF method. This process involves a background fitting to derive the ΔZ spectrum, achieved by subtracting the fitted background from the Z-spectrum value. Subsequently, the ΔZ spectrum is fitted with a line-shape characterized by three Lorentzian peaks centered at 3.5, 2.6, and 2 ppm, corresponding to amide, PCr, and Cr^+^ CEST signals, respectively. Intensities were used to report the CEST effects.

**Figure 1:**
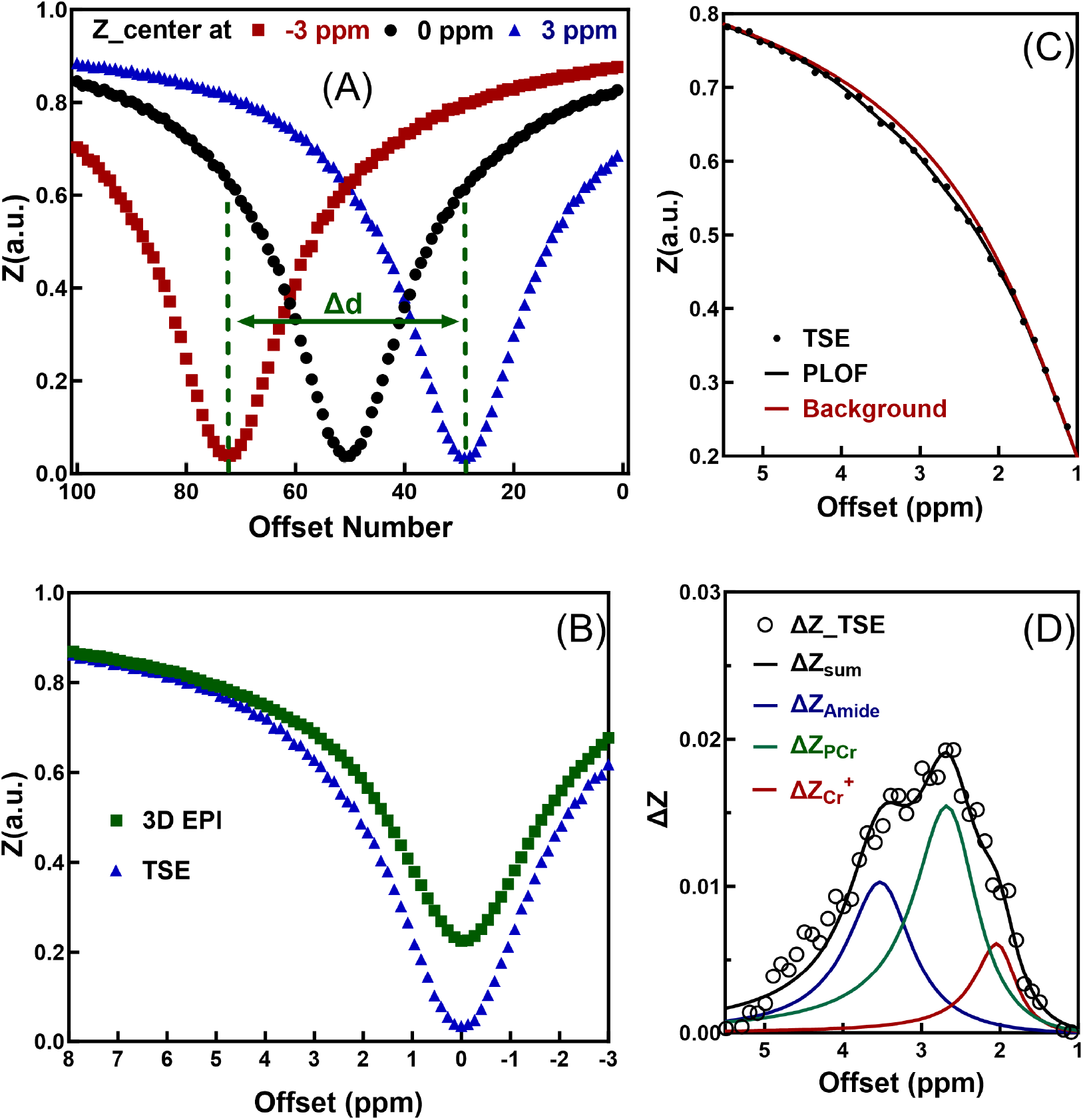
Saturation frequency calibration for UFZ spectra and extraction of CEST signals using PLOF. (A) Illustration about calibrating the saturation frequency with the spectra collected with center offsets at 3 and −3 ppm using Eqs. 6 and 7. Δd is 6 ppm. (B) Offset calibrated Z-spectra for 3DEPI and TSE readout schemes. (C) Illustration of the PLOF fitting approach with the PLOF background shown in red solid line. (D) Typical differential Z-spectrum (ΔZ) and the PLOF fitting results including amide (blue line), PCr (phosphocreatine, green line) and Cr^+^ (creatine and partial phosphocreatine, red line) CEST peaks and the sum of the three peaks (black line).

To assess the quality of CEST maps obtained with different readout schemes, typical S_0_ images at 200 ppm and three CEST contrast maps extracted via PLOF method are presented in Fig. 2 from the same subject. Regarding the S_0_ images, 3D EPI demonstrated better image quality compared to the TSE method. The blur observed in TSE images results from the extended acquisition train. Upon reviewing the three CEST contrast maps, 3D EPI exhibited greater variation compared to those obtained through the TSE method. Specifically, 3D EPI displays poorer mapping qualities with numerous values approaching zero values across all three CEST maps. Conversely, the CEST maps derived from TSE readout appear more uniform and feature higher average CEST values.

**Figure 2:**
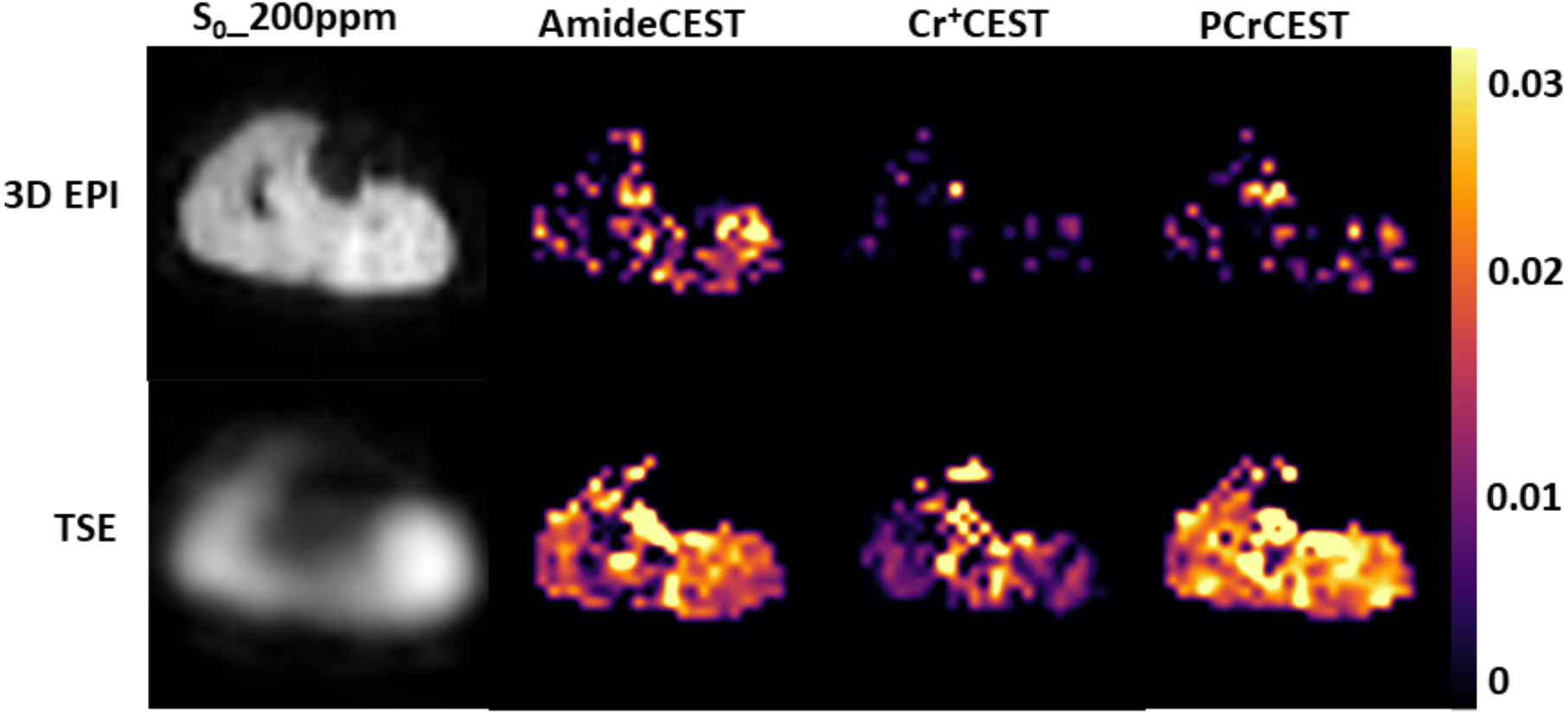
Typical S_0_ and CEST maps (amideCEST, Cr^+^CEST and PCrCEST) acquired using UFZ MRI with 3DEPI and TSE readout schemes. The CEST contrast maps were extracted with the PLOF method.

### OXPHOS measurement by PFE

To assess mitochondrial OXPHOS, the UFZ TSE method was employed with in-magnet PFE. Typical time series of CEST maps, both pre- and post-exercise, are depicted in Fig. 3. These maps encompass both the entire leg and the active muscle region (gastrocnemius), allowing for comprehensive analysis. Before the PFE exercise, the whole leg CEST maps exhibited good repeatability. However, shortly after the exercise, all maps showcased increased variation. For clarity in illustrating changes in amide, PCr and Cr^+^ signals, only the activated muscle region (gastrocnemius) was plotted separately at the bottom of the figure. Analysis of the active region revealed more pronounced CEST contrast changes post-exercise than for the whole leg. Specifically, both amide and PCr decreased after PFE and exhibited a similar increasing trend, post-exercise while Cr^+^ elevated after PFE and demonstrated a decrease during the recovery stage.

**Figure 3:**
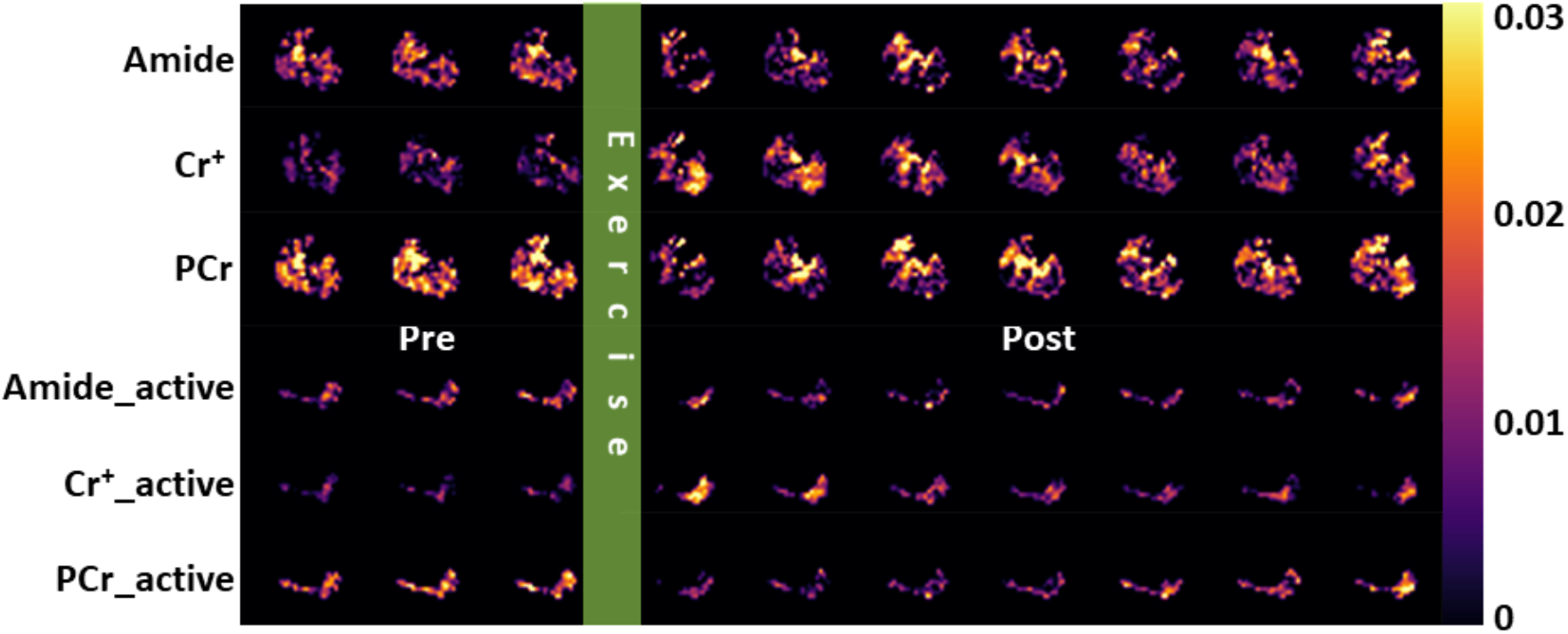
Typical whole-leg and exercise-active muscle region (gastrocnemius) CEST maps were generated both pre and post in-magnet PFE using UFZ acquisition with TSE readout. Three specific CEST maps, amide, Cr^+^, and PCr, were derived using the PLOF method. The acquisition time for each CEST map was 48.6 seconds. Both amide and PCr levels in the gastrocnemius muscle decreased following PFE, with a subsequent increasing trend noted during the post-exercise period. Conversely, Cr^+^ levels were elevated immediately after PFE and showed a decrease during the post-exercise recovery phase.

To determine the muscle-specific *τ*_*PCr*_ and 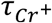 we plotted the time-series data for amide, PCr, and Cr^+^ CEST signals for the whole gastrocnemius muscle (n=9) in Fig. 4A. Additionally, the averaged time-series PCAR or CAR values (n=9) are included for comparison in Fig. 4B. Due to the extremely low signal-to-noise ratio (SNR), we opted to plot only the whole ROI for the gastrocnemius muscle. To enhance SNR, we used the averaged two PFE dynamic curves for the *τ*_*PCr*_ or 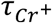 extraction on the same subject. The observed changes in amide, PCr, and Cr^+^ signals following PFE align with the trends observed in the maps (Fig. 3). The decrease in amideCEST confirms the acidic environment post intense anaerobic PFE. Subsequent pH correction using amideCEST and separate PCr/CrCEST, as per Eq. 4, reveals significantly stronger signal changes in CAR compared to Cr^+^CEST with p value of 0.0027. Based on the CEST values directly obtained from the experiments, *τ* is calculated as 87.1 ± 15.8 s for Cr^+^ and 98.1 ± 20.4 s for PCr. Following pH correction, *τ*_*CAR*_ is determined as 36.4 ± 18.6 s for Cr, while *τ*_*PCAR*_ is calculated as 43.0 ± 13.0 s for PCr.

**Figure 4:**
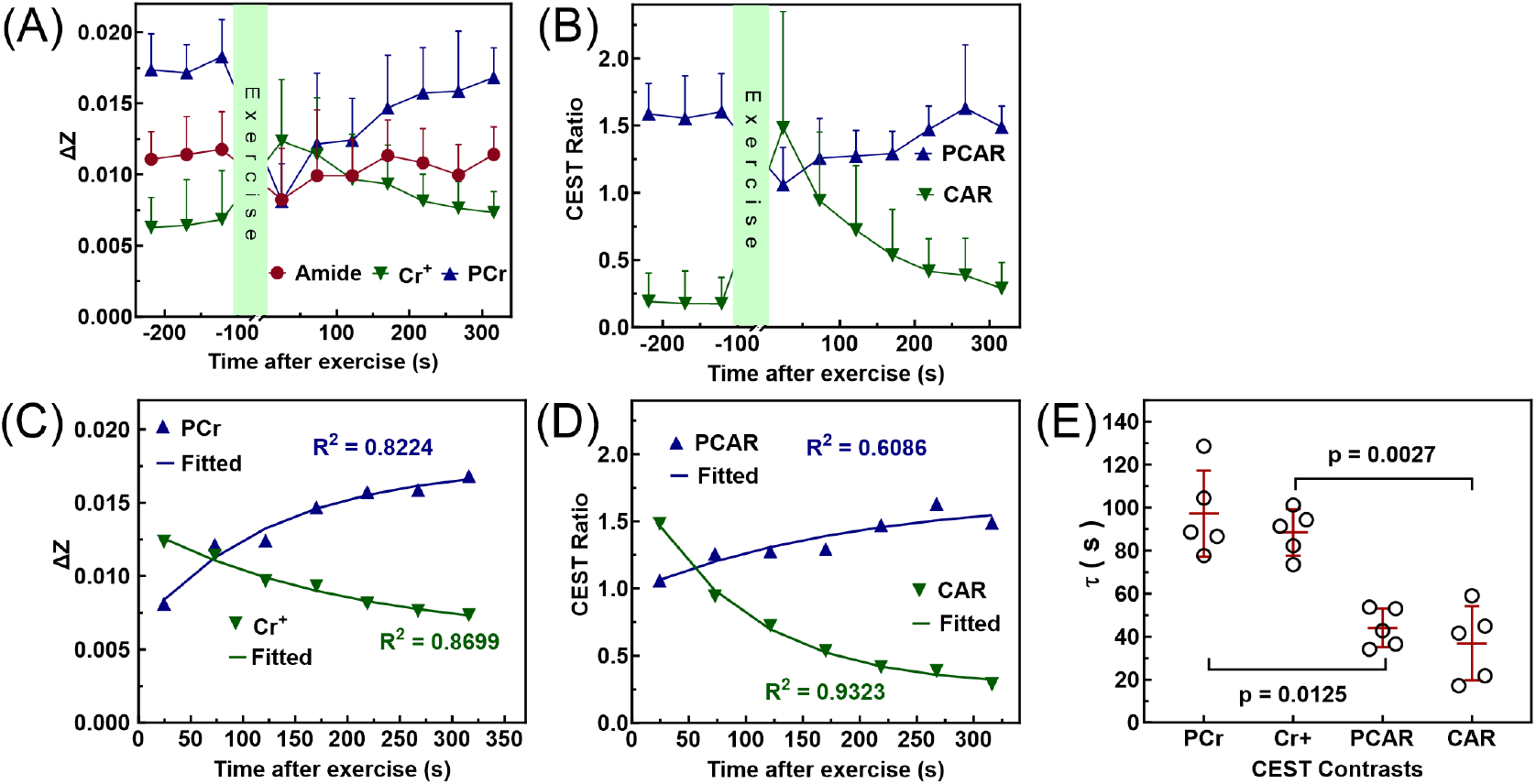
Dynamics of PCr, Cr^+^ and amide CEST signals during and post PFE. (A) Averaged amide, Cr^+^, and PCr CEST (n = 9) signal changes over time. Data from nine PFE dynamic curves across five subjects were consolidated. Consistent levels of amide, Cr^+^, and PCr are observed pre-exercise. Following FPE, a decline in amide and PCr is seen, followed by an exponential recovery. In contrast, Cr^+^ levels rise post-exercise and then diminish exponentially during recovery. (B) Ratios of PCr to amide CEST (PCAR) and Cr to amide CEST (CAR) for pH corrected CEST demonstrate clearer trends over time. (C) The fitting for the Cr^+^ and PCr dynamic curves using Equation 5 to obtain *τ*_*PCr*_ and 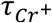 values. The R^2^ values for the fitting of averaged curves are provided, indicating a reasonable fit. (D) The fitting for the PCAR and CAR dynamic curves. (E) Compilation of *τ* values from dynamic curve fitting of PCr, Cr^+^, PCAR, and CAR for all subjects (n=5, p values were calculated by paired t test).

## Discussion

We explored the potential application of the UFZ MRI method as a rapid acquisition approach, specifically for assessing mitochondrial OXPHOS in human skeletal muscle at 3T. This was achieved by leveraging the PCr and Cr CEST signals. Given the considerable overlap of amide, PCr, and Cr^+^ CEST signals, it was crucial to acquire Z-spectra with high spectral resolution and a wide offset range to reliably extract these three CEST contrasts. Conventional CEST methods typically entail a lengthy experimental time, often spanning several minutes, due to the need for comprehensive Z-spectrum collection. However, our study demonstrated that UFZ imaging can achieve a temporal resolution of 48.6 s with reasonable spatial and spectral resolution. This advancement renders the measurement of mitochondrial OXPHOS feasible in human muscle, enabling OXPHOS function assessment for each individual muscle group using CEST MRI. In comparison, traditional ^31^P MRS typically confines measurements to averaged values of all muscle groups within the coil coverage.

Nevertheless, despite its advantages, the UFZ approach does have two significant limitations. Firstly, in achieving high spectral resolution, it sacrifices SNR. Each offset in UFZ has an effective slice thickness of only 0.4 mm, significantly thinner compared to the typical 5-10 mm slice thickness in conventional CEST. Consequently, UFZ is limited to obtaining OXPHOS function for only large ROIs. Secondly, the slice direction is utilized for encoding the offset information in UFZ. This aspect makes UFZ ideal for tissues with uniform structure, such as muscle bundles, but less suitable for tissues with high spatial heterogeneity in the slice direction.

The present study also highlighted the significant impact of acidification during PFE on PCr/Cr CEST dynamic curves. The *τ* values post PFE (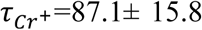 s; *τ*_*PCr*_ =98.1 ± 20.4 s) were markedly lower after pH correction (*τ*_*CAR*_ =36.4 ± 18.6 s; *τ*_*PCAR*_ = 43.0 ± 13.0 s). Notably, in ^31^P MRS with a similar PFE setup, the *τ*_*PCr*_ was approximately 30-45 s for healthy subjects (34-37), closely resembling the values obtained via PCAR. This underscores the critical importance of pH correction for reliably extracting *τ* values.

It’s worth mentioning that the amide proton transfer (APT) signal was initially proposed for tissue pH measurement (50,51). However, recent studies based on model proteins suggest that the CEST signal at 3.5 ppm, often referred to as APT signal, comprises both a pH-sensitive component, amide CEST, and a pH-insensitive component at physiological pH, amide Nuclear Overhauser Effect (amideNOE). (52,53). This dual-component nature is further supported by findings from tissue homogenates (47-51) and Z-spectral analysis in animal stroke models (54-59). The peak, termed amideCEST in this paper, originates from unstructured proteins (60,61). When utilizing the amideCEST signal for pH correction, it’s crucial to suppress the pH-insensitive amideNOE component. Recent studies on skeletal muscle at 3T, involving euthanasia of mice within the magnet, indicated effective suppression of the amideNOE components using the current PLOF approach (33).

It is feasible to enhance time resolution through the utilization of advanced acquisition approaches. However, it’s crucial to note that a GRE-type readout is not suitable for UFZ experiments. The extended acquisition train associated with GRE (1.46 s versus 0.53 s for SE-based TSE) leads to a significant loss of CEST signal due to T_1_ recovery, as evidenced by the increased signal at 0 ppm (Fig. 1B) for 3DEPI. The temporal resolution of UFZ MRI can be enhanced further using 2D MRI readout, as previously demonstrated in phantom studies (39,62). However, this approach reduces the resolution in another dimension.

## Conclusions

Our results demonstrate that UFZ MRI combined with PLOF analysis can be used for the non-invasive assessment of mitochondrial OXPHOS in the human muscle at 3T. The UFZ MRI sequence with optimized readout scheme allowed for accurate and reliable measurement of PCr/Cr dynamics with a temporal resolution superior to conventional CEST MRI techniques. The study overcomes historical challenges in ^31^P MRS by enabling the analysis of individual muscle group responses post-exercise. Notably, pH correction with amideCEST was needed to ensure the accuracy of mitochondrial function measurements, with our corrected PCr/Cr recovery rates closely mirroring those obtained via ^31^P MRS. This first demonstration of UFZ MRI in human skeletal muscle paves the way for its adoption in clinical settings, promising a new method for the evaluation of muscular health and function.

## Acknowledgments

This work was supported by P41EB031771, R01HL149742, R01AG080104, R01HL61912, and R01AG063661.

